# Benchmarking Cell Type and Gene Set Annotation by Large Language Models with AnnDictionary

**DOI:** 10.1101/2024.10.10.617605

**Authors:** George Crowley, Tabula Sapiens Consortium, Stephen R. Quake

**Affiliations:** Department of Bioengineering, Stanford University, Stanford, California, USA; Chan Zuckerberg BioHub, San Francisco, CA, USA; Department of Applied Physics, Stanford University, Stanford, CA, USA; Chan Zuckerberg Initiative, Redwood City, CA, USA

## Abstract

We developed an open-source package called AnnDictionary (https://github.com/ggit12/anndictionary/) to facilitate the parallel, independent analysis of multiple anndata. AnnDictionary is built on top of LangChain and AnnData and supports all common large language model (LLM) providers. AnnDictionary only requires 1 line of code to configure or switch the LLM backend and it contains numerous multithreading optimizations to support the analysis of many anndata and large anndata. We used AnnDictionary to perform the first benchmarking study of all major LLMs at de novo cell-type annotation. LLMs varied greatly in absolute agreement with manual annotation based on model size. Inter-LLM agreement also varied with model size. We find that LLM annotation of most major cell types to be more than 80-90% accurate, and will maintain a leaderboard of LLM cell type annotation at https://singlecellgpt.com/celltype-annotation-leaderboard. Furthermore, we benchmarked these LLMs at functional annotation of gene sets, and found that Claude 3.5 Sonnet recovers close matches of functional gene set annotations in over 80% of test sets.

## Introduction

Single cell transcriptomic sequencing (scRNA-seq) analysis has enabled discovery of novel cancer targets, rare cell types, and deepened our understanding of cell phenotype and function.^1–3^ One of the largest bottlenecks in scRNA-seq is the annotation of cell type. Until recently, this step has required input from human experts. Large language models (LLMs) have emerged as a promising tool to automate single cell analysis based on marker genes.^4^ What’s more, LLMs have shown satisfactory agreement with classical biological inference tools (i.e. Gene Ontology term analysis), and so additionally hold promise for automating interpretation downstream of cell type annotation.^5^

LLMs are primarily accessed through a commercial provider via a provider-specific interface. While some are open source and can be downloaded for local use, the size and complexity of doing so can be restrictive. Therefore, we built AnnDictionary (https://github.com/ggit12/anndictionary/), an LLM- provider-agnostic python package built on top of AnnData and LangChain that can use any available LLM by changing just one line of code (e.g. any model provided by OpenAI, Anthropic, Google, Meta, or available on Amazon Bedrock). The aim of this package is to consolidate both automated cell type annotation and biological process inference into a single python package that interfaced natively with Scanpy. Furthermore, while handling smaller datasets with ease, AnnDictionary includes optimizations to allow the LLM-based annotation of atlas-scale data. Previous work indicates that LLMs can reliably identify cell type from curated lists of marker genes—those identified via literature or calculated from previously identified cells of known type—but there has not yet been an assessment of LLMs at de novo cell type annotation, meaning annotation of gene lists derived directly from unsupervised clustering.^4^ These gene lists crucially differ from curated gene lists, because they contain unknown signal and noise that may affect the annotation process.^6,7^ De novo annotation is therefore a potentially more challenging task. The goal of this investigation is to assess the effectiveness of LLMs in this context. So, we used AnnDictionary to benchmark the de novo cell type annotation ability of the major commercially available LLMs (i.e. all models from OpenAI, Anthropic, Google, Meta, and all additional available text generation models on Amazon Bedrock, including Mistral, Titan, and Cohere). These benchmarks will be displayed as a running leaderboard at https://singlecellgpt.com/celltype-annotation-leaderboard. We also benchmarked these LLMs at functional annotation of gene sets.

Additionally, as scRNA-seq experiments continuously increase in size and complexity, so do their analyses.^8^ For example, it can be desirable to handle a dataset stratified at the donor, tissue, or cell type level, and independently operate on each of these groups of the data (i.e. normalize each tissue in Tabula Sapiens independently). AnnDictionary aims to create a formal backend for independent processing of multiple anndata in parallel. This functionality is a key building block in various scRNA-seq and spatial transcriptomic tasks (e.g. de novo annotation of cell type, label transfer, data integration, cell segmentation), and we expect AnnDictionary to serve as a useful, flexible backend for their analyses. We therefore provide notebooks of how AnnDictionary can simplify implementation of these tasks.

## Results

### AnnDictionary is a parallel backend for processing anndata

Our first step in benchmarking LLMs involved building a backend that could handle the parallel processing of many anndata through a simplified interface. The current state of the art is to manually create a dictionary of anndata objects and loop over them. We aimed to formalize this concept by: 1) defining a class called AdataDict (i.e. a dictionary of anndatas); and 2) providing an essential workhorse method—fapply—that operates conceptually similar to R’s lapply() or python’s map(), **Figure 1A**. Fapply is multithreaded by design and incorporates error handling and retry mechanisms, allowing the atlas- scale annotation of tissue-cell types by 15 LLMs in a tractable amount of time. However, multithreading can also be turned off for non-thread-safe operations.

**Figure 1.**
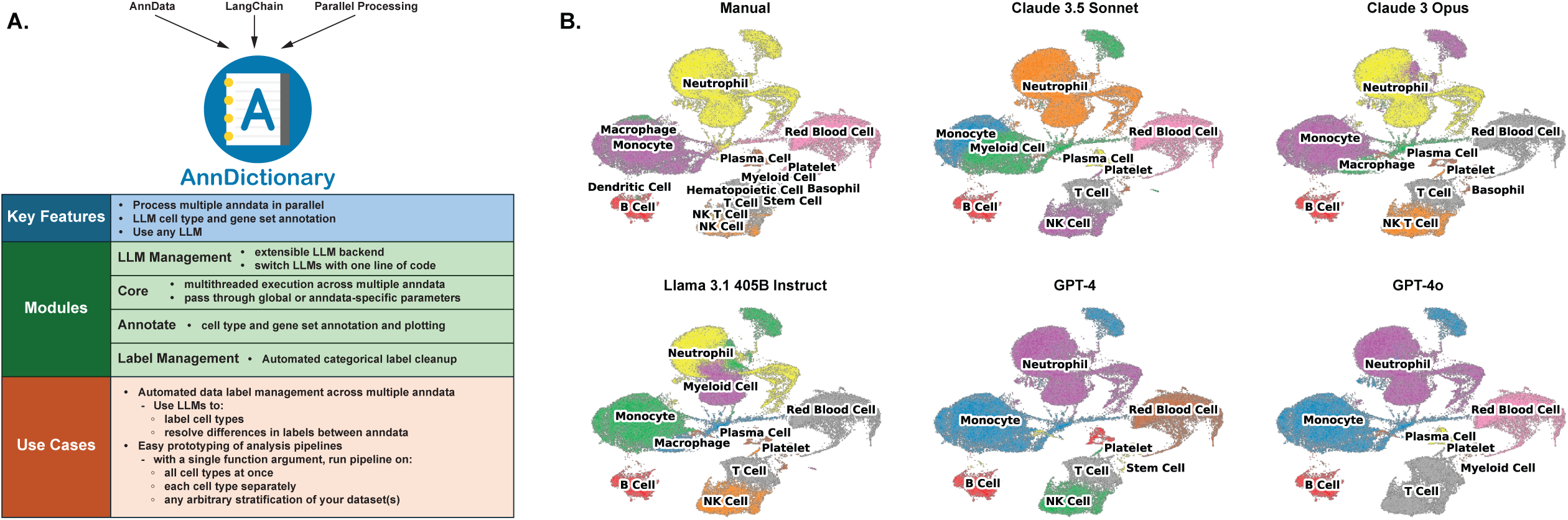
**A.** Overview of AnnDictionary—a python package built on top of LangChain and AnnData, with the goal of independently processing multiple anndata in parallel. **B.** Example LLM annotations of cell types and coarse manual annotations for all cells detected in blood of Tabula Sapiens v2.

In AnnDictionary, we also include a module of wrappers to common Scanpy functions.^9^ Furthermore, we provide centralized wrappers for label transfer pipelines (i.e. via logistic regression with Universal Cell Embedding)^10^, and data integration techniques (i.e. harmony).^11^ While this is only a small sample of available methods, AnnDictionary is extensible and can grow to accommodate additional methods. Finally, AnnDictionary functions can typically take single arguments to be broadcast to all anndata, or arguments can be provided as a dictionary, with a separate parameter for each anndata. For example, one could normalize 3 different datasets to 3 different values with a single AnnDictionary function call.

### AnnDictionary consolidates common LLM integrations under one roof

As previously mentioned, several LLM-based automations, including gene set annotation and data label management, have been developed and tested, but there does not exist, to the authors’ knowledge, a centralized implementation of them in Python built on top of AnnData, the predominant data structure used in Pythonic scRNA-seq analysis. Thus, we implemented a variety of LLM-based tools.

AnnDictionary is the first package in this space to natively support multiple LLM providers, and contains substantial technical advances over previous work,^4,12^ including few-shot prompting, retry mechanisms, rate limiters, customizable response parsing, and failure handling. All of these features contribute to a user-friendly experience when annotating datasets.

#### Cell type annotation

We designed an LLM agent that attempts to determine cluster resolution automatically from UMAP plots. Chart-based reasoning is an established task family of LLMs.^13–16^ We note that current LLMs do not seem good enough to reliably produce reasonable resolutions, but still may be a useful first pass and this capability may eventually improve. Then, we provide several functions for cell type annotation via different methods. The following methods can generally be tissue-aware at the user’s discretion. These include 1) based on a single list of marker genes, 2) by comparing several lists of marker genes using chain-of-thought reasoning, 3) by attempting to derive cell subtypes by comparing several lists of marker genes using chain-of-thought reasoning with the parent cell type supplied as context, and 4) using (2) in this list with the additional context of an expected set of cell types. At this stage, we note that, as a design principle, AnnDictionary returns relevant LLM output so that the user can manually verify cell type mappings, annotation, etc.

#### Gene set annotation

We wrote several functions to assist with gene processing. These include annotating sets of genes and adding these annotations to the metadata, for example if an “is heat shock protein” column in the gene metadata is desired. We also wrote functions that use an LLM to attempt to infer the biological process represented by a list of genes.

#### Automated label management

We implemented several functions to assist with data label management in AnnData using LLMs. Use cases include resolving syntactic differences in labels used across different studies. Some functions are built to process category labels in a single column by cleaning them, merging them, or generating multi- column label hierarchies (i.e. from cell subtype all the way to compartment). Other functions are designed to handle common situations when dealing with datasets from multiple sources or annotations from multiple methods, such as differing notation for common cell types. Furthermore, we provide ways to assess label agreement (all using LLMs to manage label comparison)

### AnnDictionary can plug in to any LLM with a single line of code

With a parallel processing backend and LLM integrations in place, the next step was to create a simple interface to allow the use of any LLM with AnnDictionary. This flexible design was desired to allow ease of use and future-proofing as new LLMs become available. To accomplish this, we built on top of LangChain to design a configurable LLM backend that we could call from the LLM integration functions without reference to a specific underlying model. The result is that the functions in AnnDictionary can be used with any LLM with just a single line of code—a function called “configure_llm_backend”.

This aspect of flexibility and incorporation of provider-specific handling, including rate limits and message formatting, enables AnnDictionary to be used to annotate Tabula Sapiens v2 with 15 different LLMs.

### Claude 3.5 Sonnet had the highest agreement with manual annotation

#### Data pre-processing, cell type annotation, and rating annotation results

For this investigation, we used the Tabula Sapiens v2 single cell transcriptomic atlas and followed common pre-processing procedures. Handling each tissue independently, we normalized, log-transformed, set high-variance genes, scaled, performed PCA, calculated the neighborhood graph, clustered with the Leiden algorithm, and computed differentially expressed genes for each cluster, see the *Methods* for details. We then used LLMs to annotate each cluster with a cell type label based on its top differentially expressed genes, and had the same LLM review its labels to merge redundancies and fix spurious verbosity. We show example LLM annotations of all cells detected in blood in **Figure 1B**.

We assessed cell type annotation agreement with manual annotation using direct string comparison, Cohen’s kappa, and two different LLM-derived ratings: one in which an LLM is asked if the automatically generated label matches the manual label and to provide a binary yes/no answer, and a second method where an LLM is asked to rate the quality of the match between automatic and manual labels as “perfect”, “partial”, or “not-matching”. Note that if the labels were a direct string match, this was treated as a “perfect” match without the need to pass to an LLM. The use of LLMs in comparing free text results is standard practice.^17,18^

To calculate Cohen’s kappa both between LLMs and with manual annotations, required a shared set of labels amongst all the annotation columns. We computed this unified set of categories using an LLM, and based calculation of all agreement metrics on these unified columns for consistency.

We ran all annotations in replicates of five to ensure stable behavior and assessment of performance, and where applicable discuss the average and standard deviation of performance across the replicates.

Claude 3.5 Sonnet had the highest binary agreement with manual annotations at 84.0 ± 0.7% of cells, followed closely by Claude 3 Opus, Llama 3.1 405B Instruct, and GPT-4o. Claude 3.5 Sonnet also had the highest binary agreement on average by cell type 70.5 ± 1.2%, and the highest proportion of perfect matches at 74.4 ± 2.7% of cells. When considering the proportion of perfect matches on average by cell type, Claude 3.5 Sonnet (54 ± 4%), Claude 3 Opus (54 ± 4%), and GPT-4o (54 ± 6%) were tied as the top performers by this metric. Finally, Claude 3.5 Sonnet had the highest percent of exact string matches across all cells (74.3 ± 2.6%), but Claude 3.5 Sonnet (53 ± 4%) was behind Claude 3 Opus (54 ± 4%), GPT- 4o (54 ± 6%), when considering the same metric averaged by cell type. Note that the performance differences among the top models are generally small, with overlapping error bounds at times. As expected, the lowest performing models were the lightweight models with smaller numbers of parameters. We omit Amazon’s Titan models from assessment because they could not reliably follow directions well enough to annotate cell types.

#### Inter-LLM agreement

The second way we assessed LLM annotation was via consistency between the LLMs. First, we measured kappa between each LLM and the manual annotation, **Table 1**. Of the LLMs tested, Claude 3.5 Sonnet Opus was the most consistent with manual annotation (κ = 0.721 ± 0.027). Then, we measured kappa values pairwise between all LLM annotation columns, **Figure 2C**, and each model’s average kappa with all other models, **Table 1**. On average, GPT-4o was the most consistent with all other LLMs (κ = 0.721 ± 0.021), and Claude 3 Opus and Claude 3.5 Sonnet were the most consistent LLM pair (κ = 0.786 ± 0.024).

**Figure 2.**
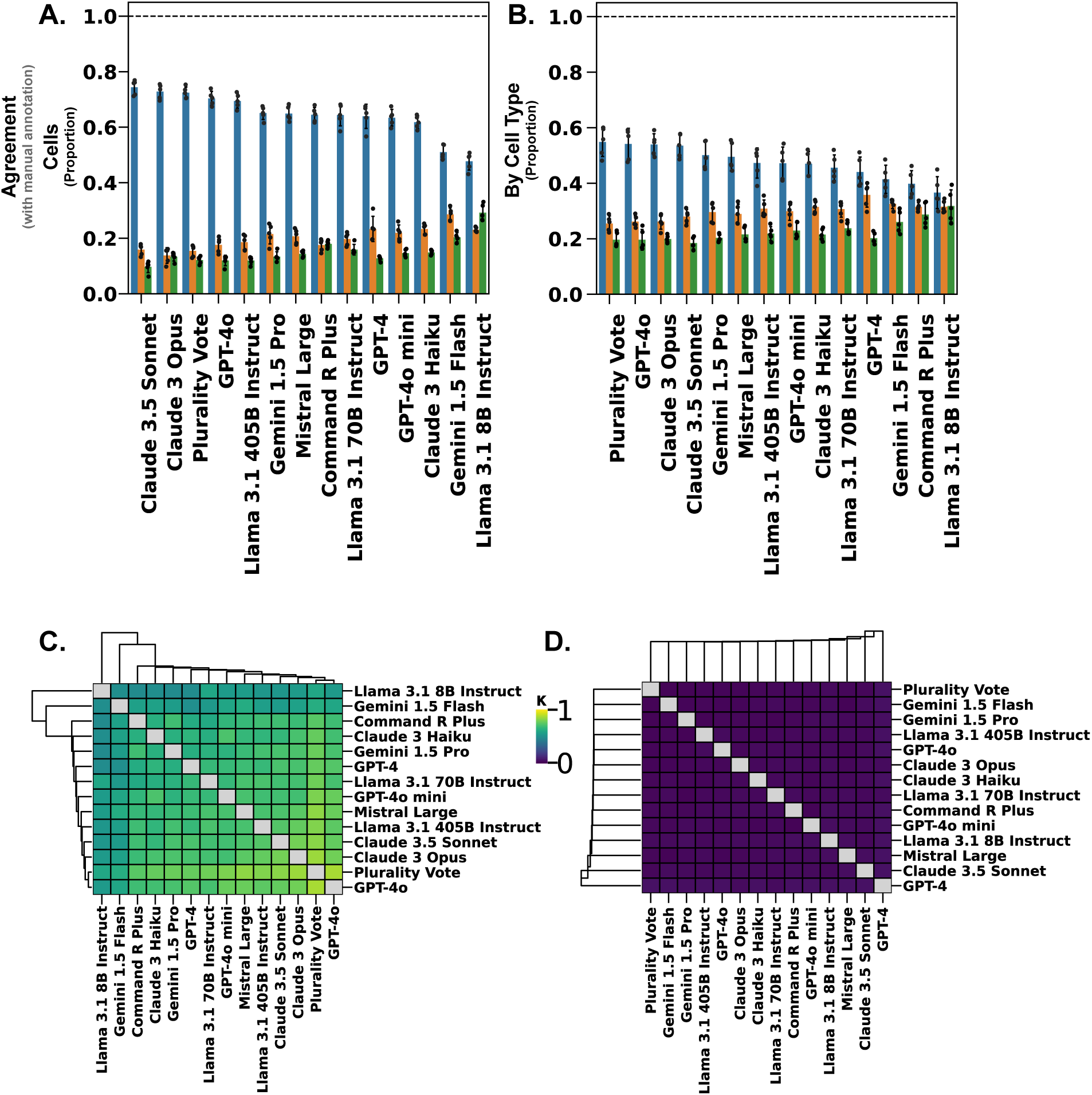
LLM cell type annotation quality as compared to manual annotation, rated by an LLM at three levels: 1) perfect, 2) partial, and 3) non-matching, and two resolutions: **A.** cells and **B.** by cell type. Inter-rater reliability measured as pairwise kappa between each LLM **C. Mean** and **D.** Standard deviation. All metrics are shown as mean and standard deviation across five replicates.

**Table 1.** LLM performance. Agreement with manual annotations measured by yes/no, quality of match, and exact string agreement. Kappa with manual annotation and average kappa of the given model with every other model. Biological process annotation of known gene lists. All values are mean ± standard deviation across five replicates.

| Model | Cell Type |  |  |  |  |  |  |  | Biological Process |
| --- | --- | --- | --- | --- | --- | --- | --- | --- | --- |
|  | Binary (%) |  | Perfect Match (%) |  | Extract String Match (%) |  | Kappa |  | Close Match |
|  | Cells | By Cell Type | Cells | By Cell Type | Cells | By Cell Type | With Manual | Average With LLMs | % of Terms |
| Claude 3 Haiku | 78.3 ± 0.8 | 66.2 ± 1.5 | 61.8 ± 2.3 | 47 ± 4 | 61.8 ± 2.3 | 47 ± 4 | 0.589 ± 0.023 | 0.652 ± 0.023 | 62.8 ± 0.4 |
| Claude 3 Opus | 82.3 ± 0.9 | 69.8 ± 1.0 | 72.8 ± 2.6 | 54 ± 4 | 72.7 ± 2.6 | 54 ± 4 | 0.704 ± 0.026 | 0.711 ± 0.020 | 71.0 ± 0.4 |
| Claude 3.5 Sonnet | 84.0 ± 0.7 | 70.5 ± 1.2 | 74.4 ± 2.7 | 54 ± 4 | 74.3 ± 2.6 | 53 ± 4 | 0.721 ± 0.027 | 0.697 ± 0.026 | 81.20 ± 0.32 |
| Command R Plus | 77.2 ± 1.0 | 59.4 ± 2.6 | 64.5 ± 2.6 | 40 ± 5 | 64.5 ± 2.6 | 40 ± 5 | 0.616 ± 0.027 | 0.646 ± 0.026 | 58.5 ± 0.7 |
| GPT-4 | 79.2 ± 0.9 | 64.3 ± 1.9 | 64 ± 4 | 44 ± 5 | 64 ± 4 | 44 ± 5 | 0.61 ± 0.05 | 0.65 ± 0.04 | 65.24 ± 0.33 |
| GPT-4o | 80.9 ± 0.7 | 70.1 ± 2.8 | 70.4 ± 2.5 | 54 ± 6 | 70.4 ± 2.5 | 54 ± 6 | 0.680 ± 0.026 | 0.721 ± 0.021 | 67.04 ± 0.33 |
| GPT-4o mini | 76.8 ± 1.0 | 66.2 ± 1.6 | 63.4 ± 3.0 | 47 ± 6 | 63.4 ± 3.0 | 47 ± 6 | 0.605 ± 0.031 | 0.681 ± 0.022 | 64.8 ± 0.5 |
| Gemini 1.5 Flash | 68.8 ± 1.3 | 60.8 ± 2.7 | 51.0 ± 2.5 | 41 ± 5 | 51.0 ± 2.5 | 41 ± 5 | 0.478 ± 0.024 | 0.561 ± 0.020 | 60.52 ± 0.18 |
| Gemini 1.5 Pro | 77.5 ± 1.8 | 67.9 ± 0.8 | 65.1 ± 2.4 | 50 ± 5 | 65.1 ± 2.4 | 50 ± 5 | 0.625 ± 0.024 | 0.658 ± 0.019 | 66.32 ± 0.11 |
| Llama 3.1 405B Instruct | 82.0 ± 1.0 | 64.9 ± 2.7 | 69.5 ± 2.6 | 47 ± 5 | 69.3 ± 2.6 | 47 ± 5 | 0.667 ± 0.027 | 0.690 ± 0.021 | 71.9 ± 0.5 |
| Llama 3.1 70B Instruct | 74 ± 4 | 61.6 ± 1.5 | 64 ± 4 | 46 ± 5 | 64 ± 4 | 45 ± 5 | 0.62 ± 0.04 | 0.665 ± 0.022 | 70.8 ± 0.5 |
| Llama 3.1 8B Instruct | 59 ± 4 | 53 ± 4 | 47.7 ± 3.2 | 37 ± 6 | 47.6 ± 3.2 | 36 ± 6 | 0.440 ± 0.031 | 0.526 ± 0.030 | 61.7 ± 0.7 |
| Mistral Large | 78.1 ± 1.5 | 66.2 ± 2.0 | 64.9 ± 2.7 | 50 ± 5 | 64.8 ± 2.8 | 49 ± 5 | 0.623 ± 0.027 | 0.696 ± 0.024 | 62.76 ± 0.17 |
| Plurality Vote | 80.5 ± 0.9 | 69.4 ± 1.7 | 72.4 ± 2.1 | 55 ± 5 | 72.3 ± 2.1 | 55 ± 5 | 0.700 ± 0.022 | 0.770 ± 0.018 | — |

#### Performance of Plurality Vote of all LLMs

We also calculated the ensemble vote of all LLMs by using the plurality label of all LLMs for each cell. In terms of performance, this method was on par with, although slightly less performant than, the other top-performing LLMs, with the exception of having the highest perfect match and exact string match rates when averaged by cell type at 55 ± 5% for both metrics, albeit by 1%.

#### LLMs excel at annotating major cell types

Overall, there is roughly a 15-20% performance difference between annotating cells vs. cell type, indicating that the models consistently agree with large, common cell types. For the subsequent sections, with the exception of **Figure 3A**, we used a single run to understand annotations by the top-performing models, and results are generally consistent between runs.

**Figure 3.**
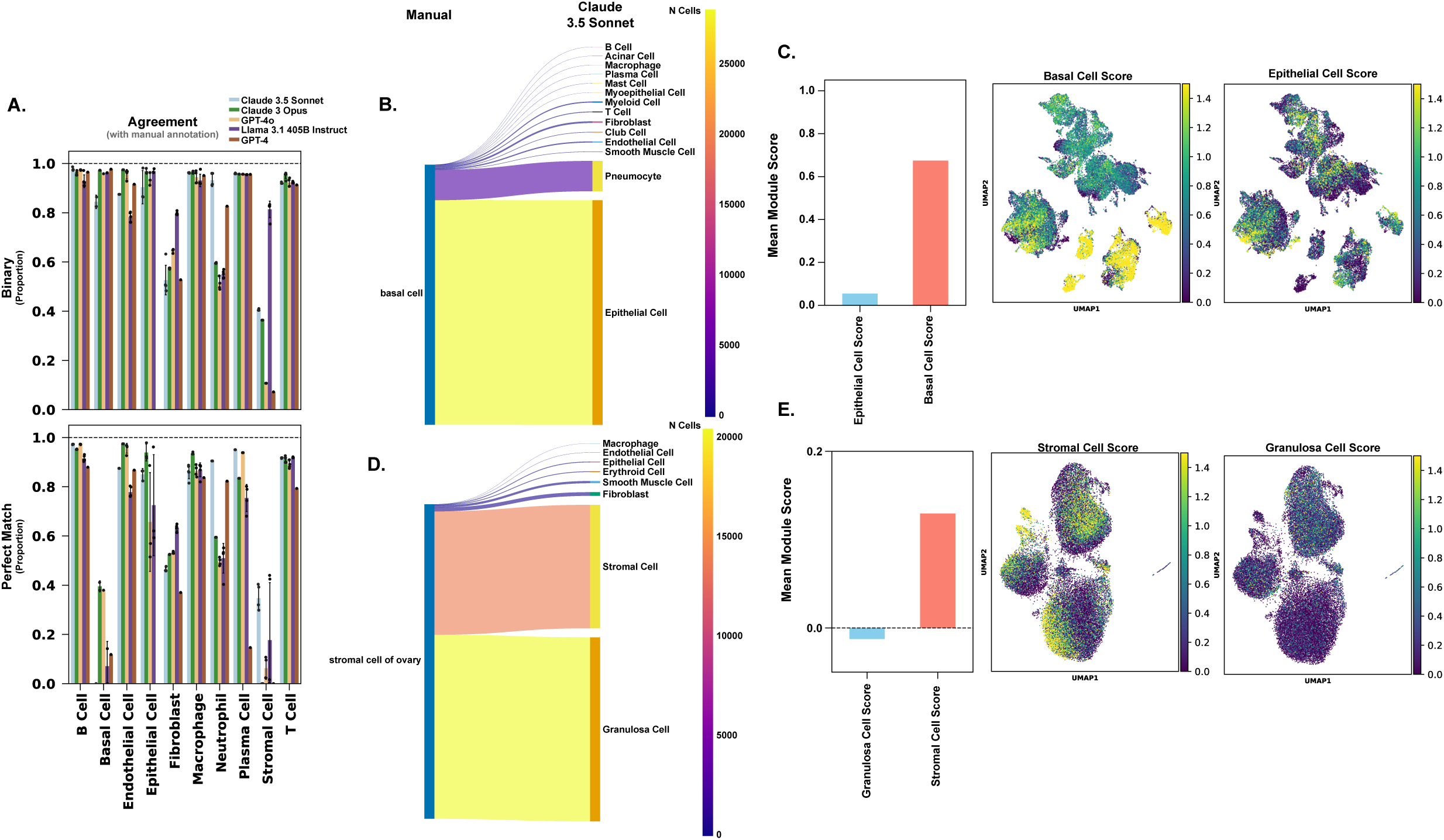
**A.** Agreement with manual annotation of top performing LLMs for the ten largest cell types by population size in Tabula Sapiens v2. As in **Figure 2**, agreement was assessed at two levels: binary (yes/no, top) and perfect match (bottom) and measured as mean and standard deviation across five replicates. For the two large cell types that disagreed with manual annotation the most: LLM annotations for cells manually annotated as **B.** basal cells and **D**. stromal cells (of ovary); and gene module scores for marker genes of the manually annotated cell type vs. marker genes for the mode LLM annotation **C.** Basal cell and Epithelial cell scores. **D.** Stromal Cell and Granulosa Cell scores.

Among the 10 largest cell types, LLMs consistently scored highly (>80-90%), except for Stromal Cells and Basal Cells, **Figure 3A**. We then looked at how the best-performing LLMs annotated these cell types. Cells that were manually annotated as basal cells were, in large part, annotated as epithelial cells by the top performing LLM (Claude 3.5 Sonnet), **Figure 3B**. Basal and epithelial cells are closely related in lineage. Based on a small number of canonical marker genes (CDH1, EPCAM, and KRT8 for epithelial cells, and KRT5, KRT14, and TP63 for stromal cells), it seems that, while there may be a subpopulation of manually annotated basal cells that have a more epithelial phenotype, basal cells dominate this group **Figure 3C**. Meanwhile, stromal cells came entirely from the ovaries, and the LLMs derived cell type names for subclusters of this population, **Figure 3D**. However, known marker genes (specifically DCN and LUM) were expressed broadly across cells manually annotated as stromal cells, **Figure 3E**.^19^ Additionally, it appeared that cells that were manually annotated as Neutrophils were consistently labeled as Macrophages by LLMs, **Supplementary Figure 2**. We also considered the LLM annotation performance within each tissue, **Supplementary Figure 3**. The tissues with the lowest average agreement between manual and LLM annotations were ear, muscle, and ovary. Viewing the tissue results in the context of the tissue-cell type level metrics, we can see that low tissue-level performance was driven by having a higher relative abundance of cell types that had low agreement with manual annotation in general. Specifically, the ear had 37% stromal cells, the muscle had 50% mesenchymal stem cells, and the ovary 72% stromal cells of ovary. So, we focus on understanding the annotations at the cell type level.

To further understand LLM annotation behavior across all cell types, we plotted inter-LLM agreement vs. agreement with manual annotation for each cell type. Among other uses, this plot was designed to allow us to identify cell types that were consistently rated by the LLMs but disagreed with the manual annotations, **Figure 4**. We separate this plot into 4 quadrants, with cell types in the 1) top-left: LLMs agree with each other, but disagree with the manual annotation; 2) bottom-left: LLMs disagree with each other and with the manual annotation; 3) bottom-right: LLMs disagree with each other but agree with the manual annotation; and 4) top-right: LLMs agree with each other and with the manual annotation. This plot is designed to qualitatively assess label confidence.

**Figure 4.**
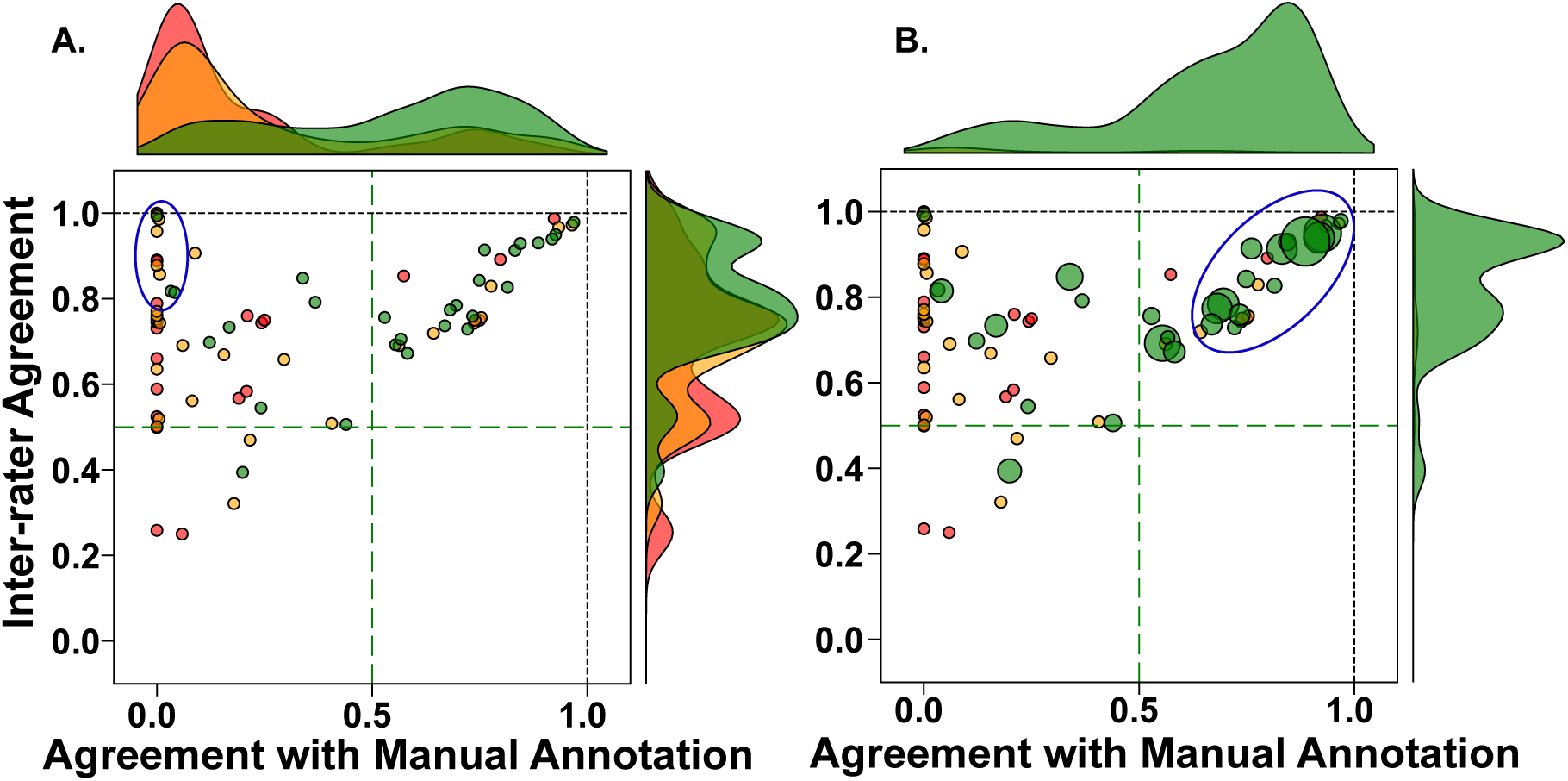
**A.** Inter-rater agreement within the top 4 performing LLMs vs. agreement with manual annotation for each manual cell type annotation, with marginal kernel density estimates stratified by tertile of cell type population size. Red, yellow, and green represent the bottom, middle, and top tertile of cell type by population size, respectively. **B.** Same set of axes as (A), with dot sizes scaled by their respective cell type populations size, and with kernel density estimates scaled by population size as well. The manually drawn ellipses outline two regions of interest: (A) the cell types with the highest inter-rater agreement and lowest agreement with manual annotation— which are the subject of **Figure 5**, and (B) the cell types with the highest inter-rate agreement and highest agreement with manual annotation—which includes the most abundant cell types discussed earlier.

Nearly all cell types had greater than 50% agreement between the LLMs, suggesting that the rate of completely spurious annotation by these LLMs is generally low across the atlas. To see how cell type population size may have affected annotation consistency, we divided cell types into tertiles of population size. Generally, major cell types had high agreement both between LLMs and with respect to manual annotation, while smaller cell types still had moderate (>50%) agreement between LLMs, but did not agree with the manual annotation. To see where the majority of cells lie, we weighted the density estimates by the number of cells in each cell type, **Figure 4B**. Here, it is clear that the majority of cells are consistently rated by the LLMs and agree with manual annotation. The differences from this trend were primarily basal and stromal cells. As previously discussed, these cells were labeled less consistently by the LLMs than most other major cell type annotations

We then investigated the group of (mostly) smaller cell types in the upper-left of this scatterplot, **Figure 4A**. For the 10 cell types that were closest to the top-left corner of the scatterplot, we plotted a confusion matrix to see the correspondence of manual annotations with annotations by the top-performing LLM (Claude 3.5 Sonnet), **Figure 5A**. The largest of these cell types were manually annotated as mononuclear phagocytes (n∼5,000), and LLM-annotated as macrophages. Based on visualization of canonical marker genes and associated module scores, there is evidence that this cluster is mostly composed of macrophages, but also contains monocytes and dendritic cells, **Figure 5B, C**. We note that there are other cell populations in the stomach that are manually labelled as monocytes and macrophages. Taken together, these data suggest that, in this case, the manual annotations may be technically correct, but the LLM annotation may be more pragmatic, as this cluster overall has highest expression of macrophage marker genes. That is to say that the label “mononuclear phagocyte” is a useful description of phenotype across several cell types, but ultimately represents a different depth of annotation than other labels in the same set. Finally, we might expect to record disagreement for small cell types whose clusters were not reproduced in the present study due to parameter decisions.

**Figure 5.**
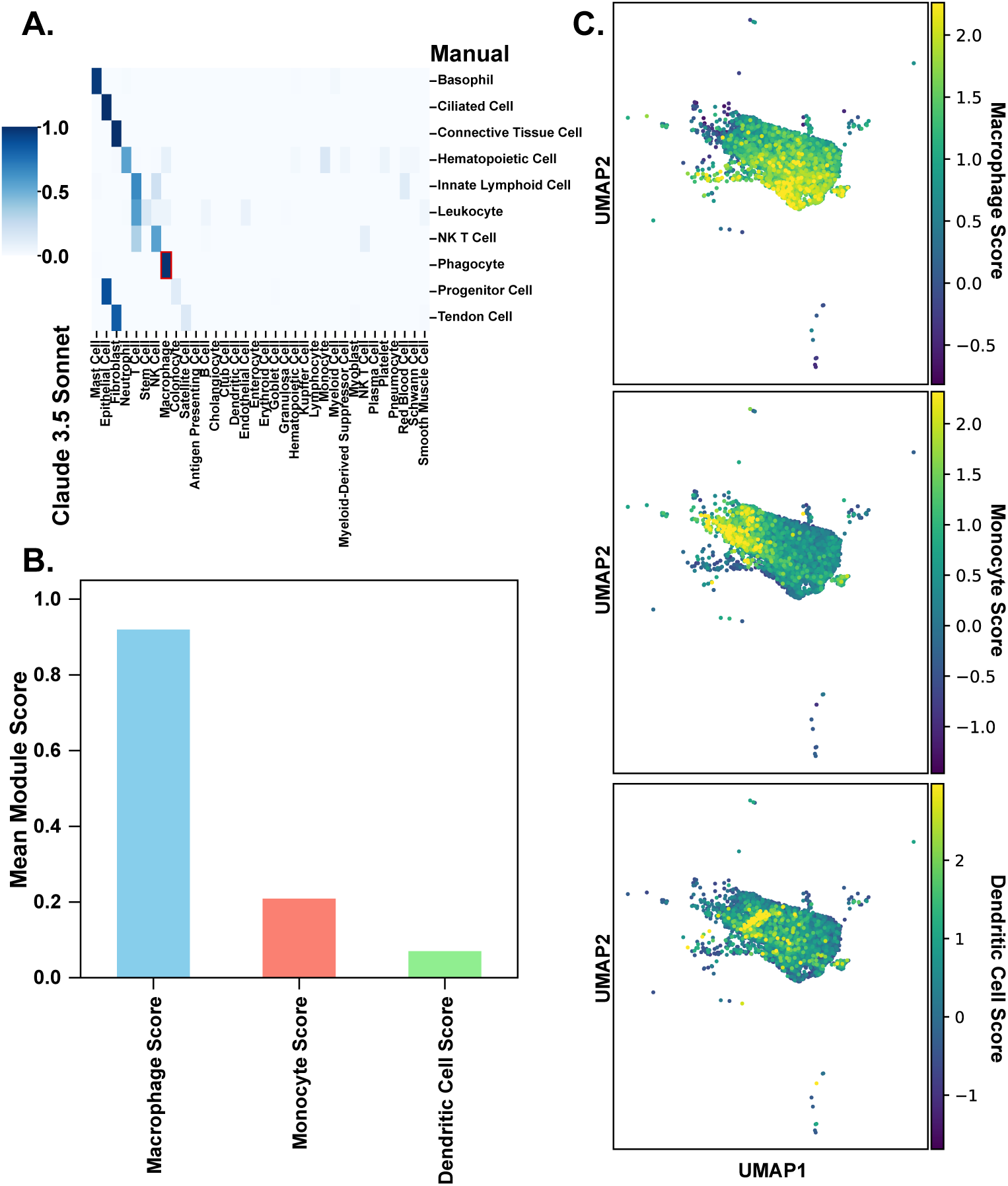
**A.** For the 10 cell types closest to the top-left corner of scatterplot in **Figure 4A**, a confusion matrix of top performing LLM annotations and corresponding manual annotations, with a red box around the largest cell type by abundance present in this group (phagocytes). The scale bar represents the proportion of cells from each category of manual annotation that are in each category of LLM annotation. Thus, each row sums to 1. **B.** Macrophage, monocyte, and dendritic cell module scores derived using canonical marker genes for cells manually annotated as phagocytes. **C.** UMAP visualization of the module scores in (B).

#### Annotation from expected cell types

We also benchmarked an LLM annotation strategy based on expected cell types. To do so, we designed a function that uses chain-of-thought reasoning to first present the list of expected cell types to the LLM, then asks it to compare and contrast the gene lists presented, then asking it to provide annotations for each list on list at a time, and finally uses partial string matching with a similarity threshold to map the LLM labels back to the initial set of expected labels. The threshold allows LLM labels that are not similar to the expected labels to pass through as new labels. Because this function uses chain of thought reasoning to help the LLM assign cell types to multiple gene lists in a single conversation, it is substantially more expensive to run and requires higher rate limits. Therefore, we benchmarked the annotation from expected cell types on a subset of the LLMs used previously that had higher API rate limits (excluding Claude 3 Opus due to cost). Here, the performances of the LLMs were only marginally lower compared to the previous annotation performances which represented a coarser scale, **Supplementary Table 1**. Furthermore, some degradation of performance is to be expected given that in this annotation strategy, the labels returned from the annotation function were not post-processed before being compared between LLMs and with manual annotations.

#### Annotation performance was not driven by the presence of cells from Tabula Sapiens v1

More than half of Tabula Sapiens v2 had not been publicly released by the knowledge cutoffs of the majority of models benchmarked in this study, including all of the top-performing models. However, cells from Tabula Sapiens v1—which was released before the knowledge cutoffs—account for a substantial portion of Tabula Sapiens v2. To determine if the presence of Tabula Sapiens v1 cells was driving the annotation results in this investigation, we re-ran the LLM benchmarking pipeline after removing cells that were part of Tabula Sapiens v1. In this control experiment, the performance of the LLMs is marginally better, **Supplementary Table 2**. This is not surprising because this test dataset was smaller. Because the Tabula Sapiens v2 dataset as a whole is of primary interest for downstream biological tasks, we still focus our investigation on the LLM annotations of the entire Tabula Sapiens v2 atlas, and use the agreement metrics calculated thereon because they are generally lower and therefore more conservative.

#### Annotation performance is robust to the LLM used in label post-processing

To address concern that the choice of post-processing model can introduce bias, we ran replicates of the post-processing step using a second LLM, GPT-4o, giving an independent assessment of the same set of annotations. We then computed the same set of performance metrics based on these corrected labels, **Supplementary Table 3**. The performance metrics as assessed by independent post-processing models are highly correlated, **Supplementary Table 4**. We thus conclude that the results are robust to choice of post-processing model.

#### Annotation performance is robust to the LLM used as rater

Self-enhancement bias refers to the potential behavior LLMs to prefer their own answers over those from other LLMs.^17^ To assess the presence of self-enhancement bias in the performance ratings presented in this study, we ran replicate experiments to compare the performances as rated by a second model, GPT-4o, **Supplementary Table 5**. The performances of each LLM when assessed by Claude 3.5 Sonnet vs. GPT-4o were highly correlated, **Supplementary Table 6**, indicating that the effect of self-enhancement bias seem to be minimal.

#### Prompt ablation study

The unablated prompt of the annotation function contained the following key components: 1) a system prompt designed to decrease output token usage and set context 2) a base prompt designed to decrease output token usage and make the LLM return only a cell type label 3) tissue context intended to provide a weak suggestion of expected labels 4) marker genes sorted as returned by scanpy’s rank_gene_groups function. To understand which components of the cell type annotation module impact annotation performance, we conducted a prompt ablation study using Claude 3.5 Sonnet, **Supplementary Table 7**. Detuning the base prompt caused annotations to be too long to use in the automated benchmarking pipeline, and so this ablation was removed from downstream comparisons. The performances amongst the remaining ablations seemed to be similar given their means and standard deviations across five replicates.

Standard deviations of metrics in the context of ablations were generally larger than in previous runs without ablation. Therefore, a dominating effect of the ablations was to decrease overall stability of the annotation pipeline. Specifically, we noticed a sharp increase in frequency of sporadic cell subtyping compared to that observed during testing of the unablated pipeline. Furthermore, without tissue context, the LLM tended to include spurious tissue information or cell types that would not reasonably be expected in the given sample. Being more stringent measures of agreement, the percent of labels that were rated as perfect matches, as well the percent of labels that were exact string matches, were more sensitive to the ablations overall than the less stringent binary rating.

#### Benchmarking biological process annotation

To assess the performance of the LLM biological process annotation function in AnnDictionary, we followed previous methodology outlined in Hu et al^5^ to define close matches between LLM-generated annotations of gene sets and the Gene Ontology terms labels from which the genes were taken, see **Methods**. All but one of the LLMs in this study—GPT-4—have not been previously benchmarked at this task. Based on annotation of 500 gene sets derived from GO Biological Process terms, Claude 3.5 Sonnet achieved the highest proportion of close matches to source GO terms (81.20 ± 0.32%), followed by Llama 3.1 405B Instruct (71.9 ± 0.5%), and Claude 3 Opus (71.0 ± 0.4%), **Table 1**.

To demonstrate the biological process annotation function, we present sample LLM-annotations of gene lists of known processes from the Gene Ontology Biological Process database. In these three cases, the LLM annotations, while slightly broader, generally agreed with the existing labels, **Supplementary Table 8**.

## Discussion

This study represents the first comprehensive benchmark of LLMs at de novo cell type annotation, and we plan to maintain a leaderboard of LLMs at this task as measured on Tabula Sapiens. We also present the first benchmarks of 14 LLMs at biological process annotation from gene sets of known process. Overall, our measures of performance indicate that large LLMs can provide reliable de novo cell type annotations at the broad cell type level, and reliable biological process annotation.

Previous work has assessed GPT-4’s ability to annotate curated lists of genes, including lists derived from Tabula Sapiens v1^4^ We build on this previous work by assessing LLMs’ abilities to annotate the full complexity of gene lists derived from unsupervised clustering. In the present study, we used Tabula Sapiens v2—which contains more than double the number of cells compared to Tabula Sapiens v1. On this larger dataset, we find the annotation of major cell types to be more than 80-90% accurate, making LLM-based annotation a viable option for first-pass cell type annotation. The flexibility of LLM-generated annotations solves a major problem of automated annotation procedures, which have historically lacked flexibility due to the need to use a reference set of annotations.^7^ Furthermore, the LLM-based approach is reference-free and so does not require additional datasets, which could otherwise increase computational burden.

In addition to direct LLM labeling of single lists of differentially expressed genes, we also tested two other annotation strategies: annotation by the ensemble vote of several LLMs, and annotation by chain- of-thought reasoning of multiple marker gene lists in a single conversation with expected cell types for context. Both of these methods add substantial runtime, financial expense, and complexity, but did yield general performance increases. Therefore, of the three methods tested, the straightforward annotation of a single gene list at a time was the best-performing, simplest, and most cost-effective approach to annotating broad cell type. Previous investigation involving curated marker gene lists found that annotations based on the top 10 differentially expressed genes gave better performance than using the top 20 or 30.^4^ Here, we used 10 marker genes and find that this gives satisfactory performance while keeping token usage minimal. Furthermore, using longer lists of marker genes risks compromising the gene lists’ specificity and including genes with smaller effect size.

Beyond saving time, effort, and cost, one major advantage of using LLMs to annotate cell types is that the LLMs seem to be able to annotate at a more consistent depth than achieved manually. However, large-scale cell type annotation with LLMs highlighted the potential pitfalls of cell type annotation in general. The apparent shortfall of LLM annotation for cell types such as basal cells and mononuclear phagocytes, may actually represent an artifact of dichotomizing continuous expressional gradients in transcriptomic data, and not speak to the performance of LLMs themselves. This is supported by the fact that the LLM annotations for these cells were, for the most part *nearly* correct, representing nearby points in the lineage of each of these cell types (i.e. a large portion of basal cells were annotated as epithelial cells and mononuclear phagocytes were annotated as macrophages). In contrast, the case of stromal cell subcluster annotation by LLMs could be an artefact of potential over-clustering in the preprocessing pipeline used in the present study, or limitations of the marker gene selection method. We also observed the difficulty inherent in distinguishing immune cell types, such as neutrophils, macrophages, and monocytes based on a small number of differentially expressed marker genes.

In the present study, we opted not to assess intra-LLM kappa (i.e. a model’s consistency with itself upon repeated, independent prompts), because we believe this to be a trivial assessment of the temperature parameter of the model. The temperature hyperparameter controls how deterministic the LLM responses are.

In addition to comprehensive cell type annotation benchmarking, we also include biological inference functions (e.g. functional gene set annotation with biological processes) and associated benchmarking. Most of the LLMs used in this study have not been previously benchmarked at this task, with GPT-4 being the one exception. We demonstrate that some LLMs have substantially higher performance compared to the best performances previously observed with other LLMs. Specifically, we observed Claude 3.5 Sonnet to achieve just over an 80% close semantic match rate when annotating curated gene sets, whereas the previous best performance was GPT-4, with a roughly 60% close semantic match rate. It is convenient to consolidate all these annotation tasks into one package. A major limitation of current gene set enrichment analyses is their dependence on the sizes of the gene sets in the query database.^20^ The use of LLMs—which do not rely on static lists of genes—to annotate pathways is therefore a promising solution to this issue.

### Limitations

There are several potential limitations related to the LLMs used in this study. We attempted to fully characterize the extent to which data leakage could influence benchmarking by denoting all information of the previous preprint that used Tabula Sapiens v2, cataloguing the use of relevant information in the present analysis, and making note of all this information in the context of the knowledge cut-offs of all LLMs used in the study. Furthermore, we ensured that the observed performances are not due to overlap with the previously published Tabula Sapiens v1 by reproducing performance in only the portion of Tabula Sapiens v2 that was not released in Tabula Sapiens v1. Because the Tabula Sapiens v2 dataset as a whole is of primary interest for downstream biological tasks, we still focus our investigation on the LLM annotations of the entire Tabula Sapiens v2 atlas, and use the agreement metrics calculated thereon because they are generally lower and therefore more conservative.

We also ensured that performance results were not due to the specific post-processing model used by reproducing highly correlated performances with another independent post-processing models.

Finally, we showed that self-enhancement bias did not substantially influence the annotation ratings by reproducing highly correlated performances with another independent models. The lack of observed self-enhancement bias is not surprising because the cell type annotations are short strings, and so it seems unlikely that they contain substantial stylistic information. While we used an LLM to rate the quality of matches, we also present these results alongside the much stricter exact string match agreement.

One goal of the present study was to build an annotation tool that could cheaply produce accurate first- draft annotations. The identification of finer-grained cell type annotations including sub type and state is often context-specific and dependent on the practitioner. So, we focus our efforts here on benchmarking annotations at a broad level, attempting to provide accurate coarse annotations to best facilitate downstream analysis. Thus, we have not considered cell type annotation beyond the broad level, but higher resolution annotations may be investigated in future work.

With regard to the benchmarking of LLMs at biological process annotation, a key limitation is that the benchmarking was performed on curated gene sets derived directly from GO terms. These lists likely differ from experimentally derived gene lists, and so further evaluation could be investigated. Difficulty in further evaluation includes establishing ground truth interpretations of gene lists, as genes are often used in many potentially independent contexts.

## Conclusion

In summary, we developed a parallel backend that simplifies processing of several anndata at once. This package is flexible to allow users to build their own additions. We also have wrapped LLM-backend configuration and switching into a single line of code, thereby simplifying the use of LLMs for annotation tasks. Beyond the benchmarking of LLMs at cell type annotation based on marker genes described here, we plan to maintain an LLM leaderboard of this task at https://singlecellgpt.com/celltype-annotation-leaderboard.

## Methods

### Data access

The Tabula Sapiens v2 dataset was accessed through its pre-release version with the help of the Tabula Sapiens Consortium. The dataset contains n=61,806 genes n=1,136,218 cells annotated by the Tabula Sapiens Consortium, more than half of which were not publicly released until December 4^th^, 2024 (this is relevant for LLM annotation, because LLMs are trained on published marker gene data).

### Data preprocessing

The full data processing pipeline, starting from raw counts, is available in the form of a snakemake pipeline at https://github.com/ggit12/benchmark_llms.

To perform the de novo LLM-based cell type annotation of the entire atlas, we first applied standard scRNA-seq analysis on a per-tissue basis with identical parameters using AnnDictionary v3.6.2 and Scanpy v1.10.2.

We first opted to use only protein-coding genes. The list of protein-coding genes is available in the benchmark_llms pipeline under benchmark_llms/src/dat/protein_coding_genes_list.csv and was downloaded from Ensemble by searching for all protein-coding human genes in the GRCh38 reference genome. We also then ensured that a common list of abundant, uninformative genes was removed (MALAT1, NEAT1, XIST, KCNQ1OT1, RPPH1, RN7SL1, RMRP, SNHG1, MIAT, H19).

All the following steps were performed with Scanpy functions, accessed through AnnDictionary wrappers to parallelize across tissue. Starting from raw counts, we 1) normalized to 10,000 counts per cell, 2) log-transformed, 3) set high variance genes (top 2000 genes), 4) scaled each gene to zero mean and unit variance, 5) performed PCA, retaining the top 50 principal components, 6) calculated neighborhood graph 7) clustered the neighborhood graph using the Leiden algorithm with resolution=0.5, 8) calculated the UMAP embedding, 9) calculated differentially expressed genes for each cluster using the t-test and Benjamini-Hochberg-corrected p-values. All of these preprocessing parameters are within the range of those commonly used. With regard to the resolution for Leiden clustering, we opted to use the same moderate-to-low resolution value across all tissues. We initially tested higher resolutions in the range 2-5 with the intent to merge clusters based on their cell type identities. But we observed that doing so often yielded clusters that were enriched for genes typically thought of as artifactual such as mitochondrial and ribosomal signal. Thus, we caution the user against the over-cluster-and-merge strategy.

### LLM hyperparameters

For all LLM queries, we set the temperature to 0, which makes the LLM behave more deterministically. For all other hyperparameters, the defaults were used.

### Cell type annotation

We then annotated clusters with cell type labels using the following process, which contains multiple LLM passes over the labels. For the following analysis, each tissue is handled entirely separately. First, using AnnDictionary’s ai_annotate_cell_type() function, clusters within a tissue were independently annotated with an LLM based on the top 10 marker genes.^4^ Second, using AnnDictionary’s simplify_obs_column() function, cell type labels within a tissue were merged via the same LLM as used in the previous step to account for redundant labels (i.e. “Macrophage”, “macrophage”, and “macrophage.”), sporadic cell subtyping (i.e. “T cell” and “Cytotoxic T cell”), and sporadic verbosity (i.e. “This cell type looks like a Macrophage. I’m basing this off expression of…”). We performed correction for subtyping because we noticed that, during package development, broad cell type labels were easy to consistently elicit, but more specific labels were not consistent. We therefore performed agreement assessments at this coarser cell type level. This may not speak to LLM performance specifically, and could be due to conflicting or undecided literature on cell subtypes in general.

We performed this two-pass annotation process with each of the LLMs listed in **Table 2**.

**Table 2.** List of LLMs studied.

| Company | Provider | Endpoint | Knowledge Cutoff Date |
| --- | --- | --- | --- |
| Meta | Bedrock | meta.llama3-1-8b-instruct-v1:0 | December 2023<sup>21</sup> |
| Meta | Bedrock | meta.llama3-1-70b-instruct-v1:0 | December 2023<sup>21</sup> |
| Meta | Bedrock | meta.llama3-1-405b-instruct-v1:0 | December 2023<sup>21</sup> |
| Amazon | Bedrock | amazon.titan-text-express-v1 | Not defined<sup>22</sup> |
| Amazon | Bedrock | amazon.titan-text-lite-v1 | Not defined<sup>22</sup> |
| Cohere | Bedrock | cohere.command-r-plus-v1:0 | February 2023<sup>23</sup> *not clear which model version is accessed on bedrock |
| Mistral | Bedrock | mistral.mistral-large-2407-v1:0 | Released July 24<sup>th</sup>, 2024<sup>24</sup> |
| Google | Google | gemini-1.5-pro-002 | September 2024<sup>25</sup> |
| Google | Google | gemini-1.5-flash-002 | September 2024<sup>25</sup> |
| OpenAI | OpenAI | gpt-4-0613 | November 30<sup>th</sup>, 2023<sup>26</sup> |
| OpenAI | OpenAI | gpt-4o-2024-08-06 | September 30<sup>th</sup>, 2023<sup>26</sup> |
| OpenAI | OpenAI | gpt-4o-mini-2024-07-18 | September 30<sup>th</sup>, 2023<sup>26</sup> |
| Anthropic | Anthropic | claude-3-5-sonnet-20240620 | April 2024<sup>27</sup> |
| Anthropic | Anthropic | claude-3-opus-20240229 | August 2023<sup>27</sup> |
| Anthropic | Anthropic | claude-3-haiku-20240307 | August 2023<sup>27</sup> |

\* not clear which model version is accessed on bedrock

### Annotation post-processing

In order to compute inter-LLM consistency, we needed to have a shared set of labels across all LLM-generated and manual annotations. Our goal was to build an automated assessment pipeline so that the benchmarking results can be updated as new models become available. Therefore, to automate annotation post-processing, we used an LLM to create a unified set of cell type labels across all annotations (manual and LLM-based). We opted to use Claude 3.5 Sonnet for label unification because, during testing, we observed that it had the best reasoning capabilities and sufficiently high API rate limits. We also compared to results when using GPT-4o for the same task to assess if the choice of model biased the results. GPT-4o was chosen for this step due to sufficient API rate limits, similar reasoning capabilities, and having come from an independent provider compared to Claude 3.5 Sonnet.^28^

### Cell type annotation by multi-LLM vote

To investigate the extent to which the annotation performance varied by using the ensemble vote of the LLMs, we computed the consensus annotation for each cell as the plurality vote of all LLMs, and rated annotation agreement in the same manner as the other LLM annotations.

### Agreement with manual annotation

To assess label agreement between the LLMs and manual annotations, we used an LLM (Claude 3.5 Sonnet) to automate the comparison of cell type labels because this allowed us to meet the scale requirements and goals of the present study, which involved assessing the agreement of ∼10,000 unique pairs of labels. We measured label agreement by comparing LLMs to 1) manual annotation 2) each other.

#### Resolutions

We calculated the rate of agreement of each model with manual annotations at four resolutions: 1) cells, 2) cell types, **Table 1**, 3) tissue-cell types, **Supplementary Figures 1 and 2**, and tissues, **Supplementary Figure 3**. The first resolution, cells, corresponds to treating each cell individually, and seeing the rate of agreement across all cells. We consider this resolution to be the most representative of overall annotation performance, because it directly measures the overall proportion of cells that are correctly annotated across the dataset. The second resolution, cell types, is calculated by taking the mean of agreement of cells when grouped by the manual annotation column. The average of this metric across all cell types is presented in **Table 1**, and in specific cell types in Figures 3A and 3B. This metric is useful for uncovering annotation performance of the model at each cell type, but, unlike the cell resolution, does not take into account differential cell type abundance. The third resolution, tissue- cell type, is calculated as the mean agreement of within a single tissue-cell type. This metric was considered because cell types are known to have tissue-specific expression programs, and so it was possible that the resulting differentially expressed genes, and thus annotation performance, varied at the tissue-cell type level. Finally, agreement was calculated at the tissue level by taking the mean agreement of all cells in a single tissue.

#### Agreement metrics

We assessed agreement between LLM and manual annotations using several different metrics.

First, for each unique pair of manual and LLM cell type labels, we used a function that asks the LLM if the two annotations agree, and then process the response to get a binary output (0 for no, 1 for yes). Second, we asked the LLM to rate the quality of the label match as perfect, partial, or non-matching and calculated the rate of “perfect” matches among individual cells and cell types, **Table 1**, and visualized the rates of all match qualities, **Figure 2A, B**. Third, we assessed the agreement via direct string comparison between the labels.

We note that the direct string comparison is overly conservative and represents a lower bound on performance. This is because cell type labels can agree semantically, but not typographically, for example “T-helper cell” and “CD4+ lymphocyte.”, or be typographically close but semantically far, for example “T cell” and “B cell”. It would be difficult to programmatically resolve these differences. However, it is standard practice to use LLMs to compare and rate the quality of free text results.^17,18^ Our rationale for using an LLM here was that it offered a powerful solution for assessing conceptual distances between free text labels. Finally, there is a standard ontology for cells. However, the first step of annotation typically uses free text labels, and most dataset annotations are not mapped to terms in the standard ontology. Furthermore, converting free text labels to standard terminology can lose specificity desired by individual research projects for their particular applications.^29^ Annotation is also an iterative process that involves updating labels with the most up-to-date knowledge, and there can be lag in updating formal ontologies. The tool developed in this study is designed to be used in tandem with a researcher to increase the speed of draft annotation generation. Thus, we believe it is of most relevance from a practical perspective to assess the quality of free text labels provided by LLMs, as that is the type of label that will most likely be used by the majority of users, especially at early stages of projects.

Finally, we measured Cohen’s kappa between each LLM label column and the manual label column as additional metrics of agreement.

### Annotation from expected cell types

The annotation from expected cell types was carried out starting from the same differentially expressed gene lists as used in the previously mentioned annotation benchmarking. To annotate gene lists, we used the ai_annotate_from_expected_cell_types() function from AnnDictionary, passing the unique cell types present in the given tissue as the expected cell types. This function uses a chain-of-thought reasoning approach where first, the LLM is supplied the tissue and expected cell types as context in the system prompt, where supported, or a message from the user if system prompts are not supported. Then, all differentially expressed gene lists to be annotated are supplied, and the LLM is asked to compare and contrast them. Finally, each list is presented again and requested to be labeled. The last step of this function is to use partial string matching to map the LLM-supplied labels back to the expected cell types, with a similarity threshold to allow new labels from the LLM to pass through to the returned annotations. Because the returned annotations are generally mapped back to the original set of expected annotations, these annotations are directly used to calculate agreement metrics without the postprocessing step mentioned in the previous annotation pipeline (described in the Methods section titled “Annotation post processing”). The source code modifications for this analysis are available under the from_expected branch of the benchmark_llms code repository.

### Assessment of self-enhancement bias

To assess the presence of self-enhancement bias on the performance ratings presented in this study, we ran replicate experiments to compare the performances as rated by a second model from an independent provider, GPT-4o, selected for its sufficiently high API rate limits and reasoning capabilities. For consistency, the same set of post-processed annotations were used here as were used in the main assessment of performance. The source code modifications for this analysis are available under the cross_check branch of the benchmark_llms code repository.

### Inter-LLM agreement

The second way we assessed LLM annotation was via consistency between the LLMs. To do so, we measured Cohen’s kappa between each pair of LLM label columns. We also computed each LLM’s average consistency with other models as the average of pairwise kappas for that LLM.

### Qualitative assessment of label confidence

To qualitatively assess label confidence of manual annotations, we used a scatterplot where each cell type was plotted based on how consistently the LLMs annotated the cells in that cell type, and how much these LLM labels agreed with the manual annotation. To do so, we opted to use only the top (n=4) performing LLMs when ranked by their overall binary agreement (percent of cells). We chose n=4 because during testing, the top 4 LLMs tended to include models from several companies, and so we hoped to sample across a potentially more diverse array of LLM behaviors and identify annotation inconsistences that were robust to model- or company-specific behaviors.

Inter-LLM agreement was calculated as the percent of LLM labels that matched the plurality label among the top LLM models for that cell. LLM agreement with manual annotation was calculated as the percent of LLM labels that matched the manual annotation for that cell. Both inter-LLM agreement and LLM agreement with manual annotation were calculated per-cell, and averaged across all cells of a given cell type based on their manually annotated cell type. We also calculated the number of cells in each cell type, which we used to split the cell types into three groups which represented the smallest, middle, and largest cell types by population size. We calculated and plotted Gaussian kernel density estimates of cell types on each axis—inter-LLM agreement and agreement with manual annotation—and scaled the areas of the marginal density estimates by the total number of cells they represented, to understand the distribution of cell types on this plot, where the majority of cells lie on the agreement plot, and how agreement varied by cell type population size.

### Risk of data leakage biasing the results

To understand the risk of data leakage, we reviewed the release dates of all manuscripts that used Tabula Sapiens v2, and looked at these in the context of the model training knowledge cutoff dates. Prior to the present manuscript (first posted in preprint form on bioaRxiv on October 13th, 2024), the only work to use Tabula Sapiens v2 data was posted on bioaRxiv on November 29^th^, 2023. Six canonical memory vs. naïve B cell markers (IGHM, IGHD, YBX3, TNFRSF13B, CD27, ATXN1) were the only differentially expressed genes from Tabula Sapiens v2 that were mentioned. Given that these are all previously published B cell markers,^30–32^ their mention in Rosen et al. does not provide noteworthy additional information in LLM model training compared to what already existed in the literature. For completeness, we assessed the frequency of these genes in the full list of all marker genes used for annotation in this study. Of 676 clusters annotated, YBX3 was used in the annotation of 10 clusters, and none of the other 5 genes were used. Any other preprints or publications that used Tabula Sapiens v2 were released after knowledge cutoff dates of models used in the present manuscript, **Table 2**. There were 3 exceptions, which involved models that were in the lower range of performance. Specifically, Mistral Large 24.07 was released July 24^th^, 2024, but Mistral does not specify a knowledge cutoff date. The two Titan models also do not have a knowledge cutoff date, but these models were omitted from detailed benchmarking in this study as previously discussed. We thus conclude that there is minimal risk of data leakage biasing the annotation results.

### Ensuring pipeline stability

To ensure the stability of the results, we analyzed the performance of the LLMs across five replicate annotation runs and report performance metrics as mean ± standard deviation.

### Validation on Tabula Sapiens v2

Using the Tabula Sapiens v2 dataset obtained as described above in *Data access*, we took only donors that were not included in Tabula Sapiens v1, that is any donor after but not including TSP15. We then re- ran the entire LLM benchmarking pipeline on this subset of the data. We excluded pancreas from this analysis due to low cell abundance. The source code for this version of the pipeline is available in the benchmark_llms GitHub repository under the branch metrics_only.

### Prompt ablation study

We generated a set of ablated functions by taking the core annotation function (ai_cell_type) and independently ablating each component of the prompt. The first ablation was to detune the base prompt from, “In a few words and without restating any part of the question, describe the single most likely cell type represented by the marker genes:” to, “What cell type is represented by the marker genes:”. The second ablation was to remove the tissue context from the prompt. The third ablation was to remove the system prompt, “You are a terse molecular biologist.” The fourth and final ablation was to randomize the gene order in the gene list input. To test how these ablations impacted the agreement of LLM-generated annotations and manual annotations, we used claude-3-5-sonnet-20240620 on the full Tabula Sapiens v2. We followed the same data preprocessing and filtering steps as used in the main analysis, as well as the same label comparison procedure. The source code for the modifications used in the prompt ablation study are available in the benchmark_llms GitHub repository under the branch ablate.

### Benchmarking biological process annotation

To benchmark LLMs at biological process annotation, we followed the procedure designed by Hu et al.^5^ We used gseapy v1.1.8 to access the 2023 release of the Gene Ontology Biological Process (GOBP) database (n=5,406 human terms). To generate gene lists of known biological process, we randomly selected 500 GOBP terms from the set of GOBP terms with between 3 and 100 genes (n=5,094). Each of these 500 gene lists were annotated by LLMs using the ai_biological_process function from AnnDictionary. To measure whether each LLM annotation was a close semantic match to the gene list’s associated GOBP term, we computed text embeddings of all LLM generated annotations and all 5,406 GOBP terms, and used the cosine similarity to measure the similarity between the embeddings of each LLM annotation and all GOBP terms. An LLM annotation was considered a close match to the source GOBP term if the source GOBP term was in the 95^th^ percentile or higher of GOBP terms by cosine similarity of the text embedding to the LLM annotation. Text embeddings were computed using OpenAI’s text-embedding-3-large model. We ran 5 replicates of each model to ensure stability, and also tested several random seeds to ensure that observed performance was not due to the specific set of GOBP terms sampled. Source code to reproduce this benchmarking analysis is available in the benchmark_llms GitHub repository as a Jupyter notebook in the nbs directory.

Then, to make a brief example, we retrieved the gene lists associated with the first three terms in the database, and used AnnDictionary’s ai_biological_process function to label the gene sets with claude-3- 5-sonnet-20240620.

### Correlation analysis

Correlations and p-values presented in Supplementary Tables 4 and 6 were calculated using scipy v1.14.1 with the function scipy.stats.pearsonr and p-values are two-sided.

## Data Availability

This study used publicly available data. Instructions to access the dataset can be found at https://tabula-sapiens.sf.czbiohub.org/whereisthedata. Source data are provided with this paper.

## Code Availability

The code to run the LLM benchmarking pipeline and reproduce all figures and tables presented in this manuscript is available at https://github.com/ggit12/benchmark_llms. The AnnDictionary source code is available at https://github.com/ggit12/anndictionary.

## Acknowledgements

We thank Jaeyoon Lee for his insights and feedback on the python package and the manuscript. We thank Mira N. Moufarrej for her feedback on the python package. We thank Fabio Zanini, Madhav Mantri, Loïc Royer, and Yusuf Roohani for discussions related to the manuscript and python package. We thank Robert C. Jones and Jaeyoon Lee for help accessing the Tabula Sapiens v2 dataset.

