## Supplementary Figures and Tables for "Benchmarking Cell Type and Gene Set Annotation by Large Language Models with AnnDictionary"

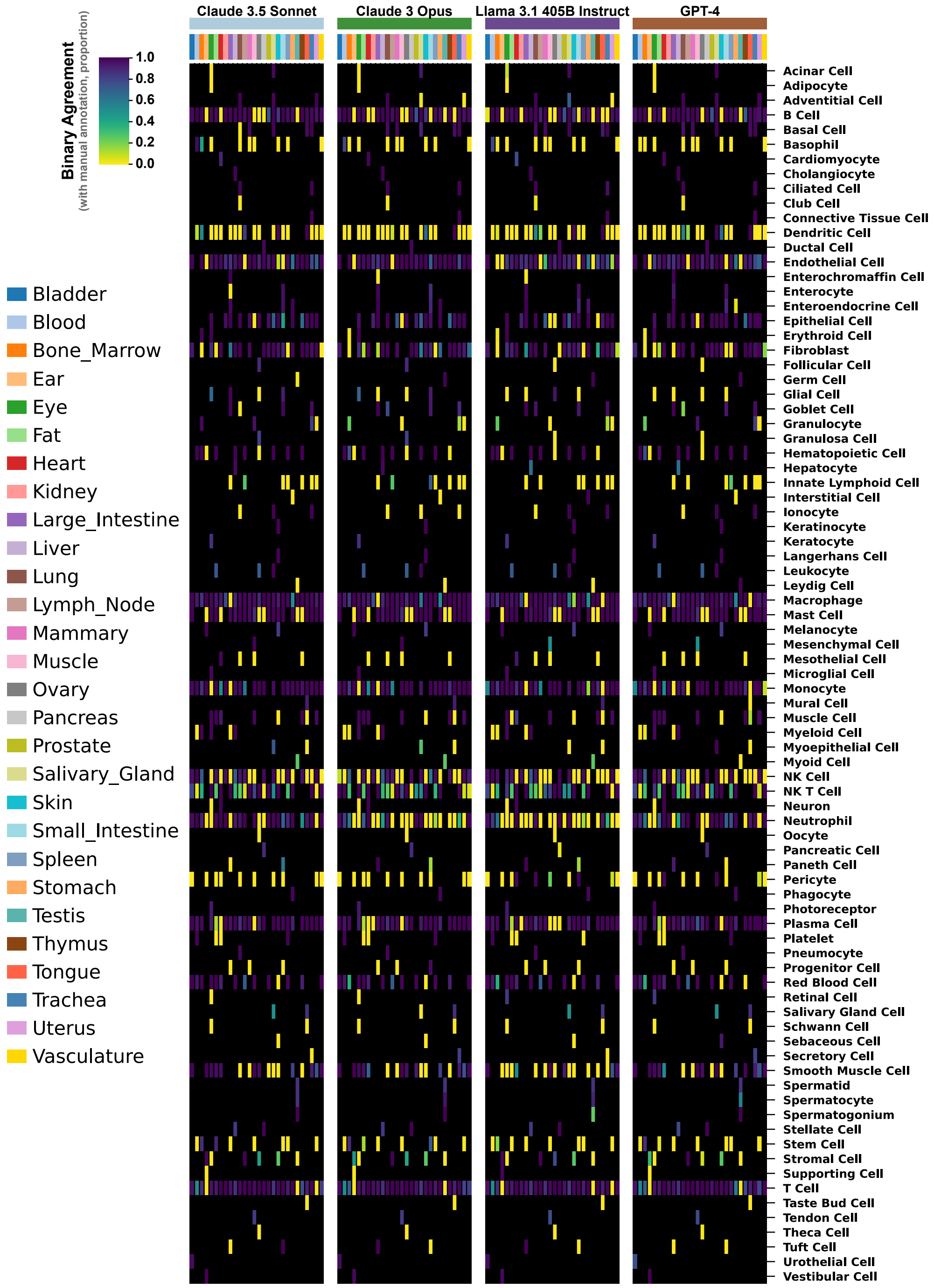


**Supplementary Figure 1.** Rates of agreement with manual annotation as measured by a binary response for the four large language models with the top performance based on this metric in this exemplar run. Data is plotted at the tissue-cell type resolution, where columns are tissues and rows are cell types, repeated for each large language model.

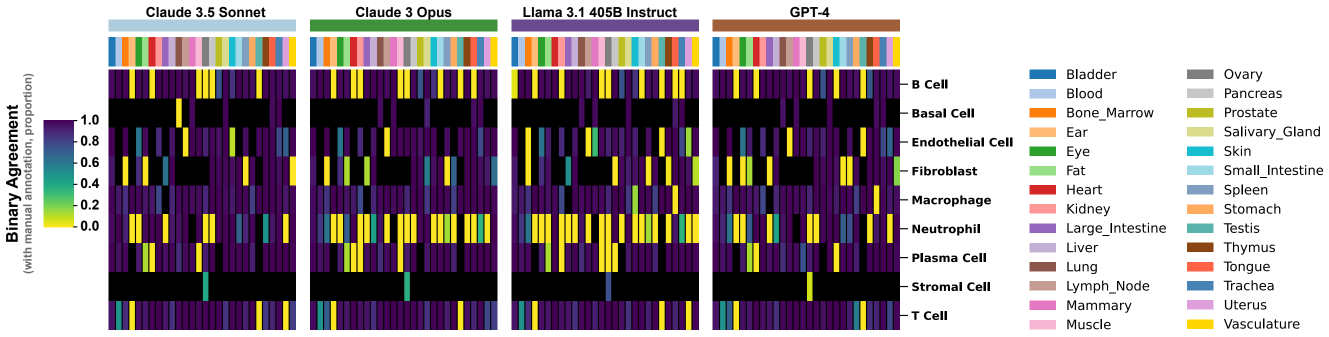


**Supplementary Figure 2.** Rates of agreement with manual annotation for the 10 largest cell types by number of cells available. As in **Supplementary Figure 1**, agreement is measured as a binary response, data is plotted at tissue-cell type resolution, and visualization is repeated for each of the four large language models with the highest overall binary agreement rate in this exemplar run.

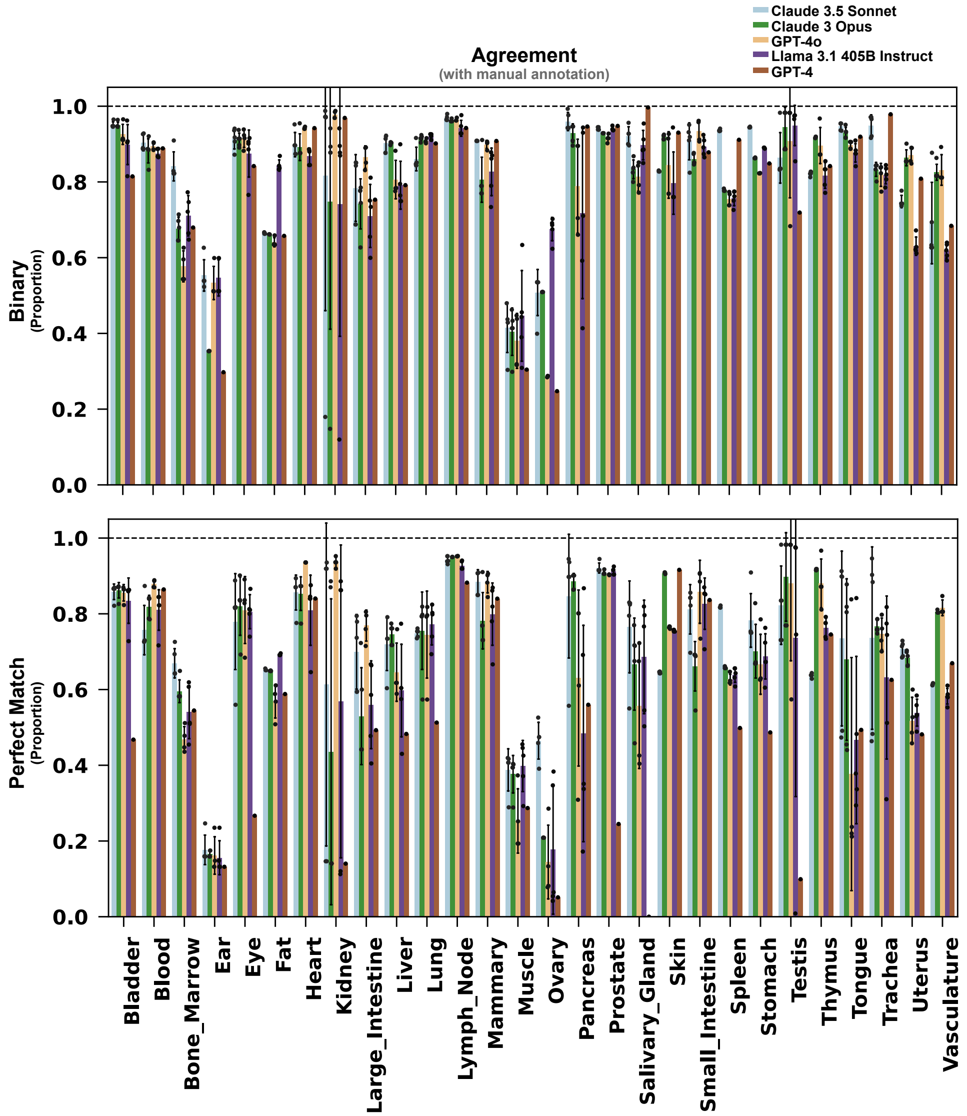


**Supplementary Figure 3.** Agreement with manual annotation of top performing LLMs for each tissue in Tabula Sapiens v2. As in **Figure 2**, agreement was assessed at two levels: binary (yes/no, top) and perfect match (bottom).

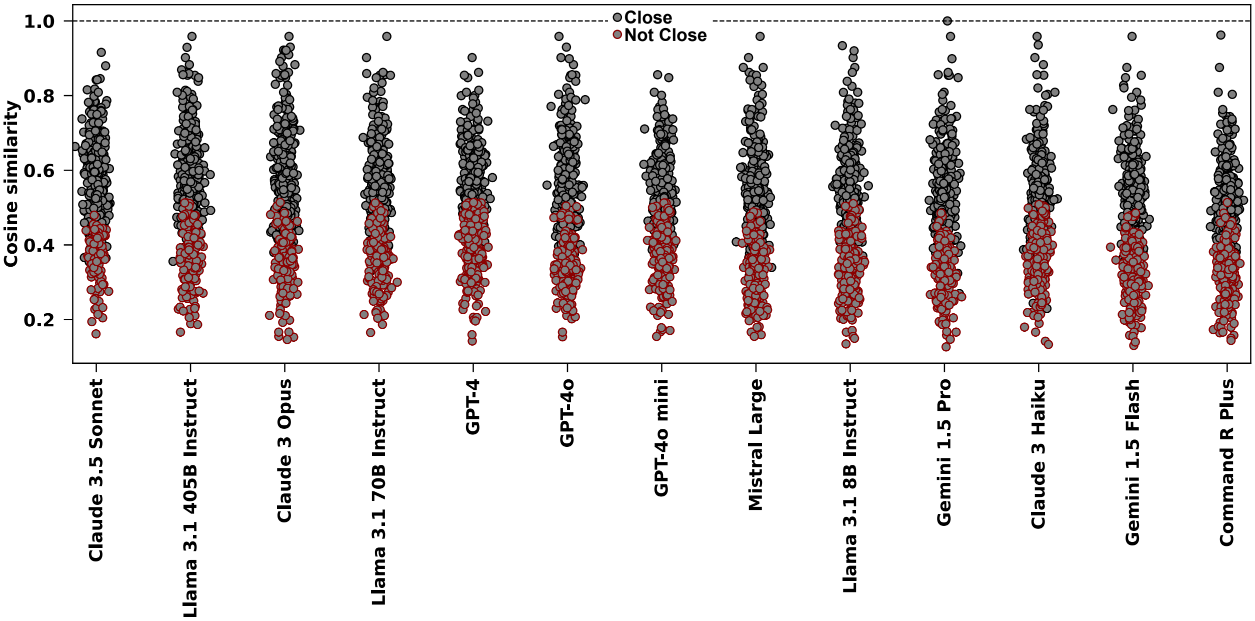


**Supplementary Figure 4.** Distribution of cosine similarities between Gene Ontology Biological Process term and LLM annotation of the genes in that term, colored by whether the LLM annotation was a close match to the source GO term as defined in the Methods, shown for one exemplar run.

**Supplementary Table 1.** LLM performance when annotating from a set of expected cell types using chain-of-thought reasoning. Agreement with manual annotations measured by yes/no, quality of match, and exact string agreement. Kappa with manual annotation and average kappa of the given model with every other model. All values are mean ± standard deviation across five replicates.

| | Binary (%) | | Perfect Match (%) | | Extract String Match (%) | | Kappa | |
|---|---|---|---|---|---|---|---|---|
| | **Cells** | **By Cell Type** | **Cells** | **By Cell Type** | **Cells** | **By Cell Type** | **With Manual** | **Average With LLMs** |
| **Claude 3 Haiku** | 71.9 ± 1.6 | 45.4 ± 0.8 | 45.5 ± 1.3 | 25.2 ± 0.7 | 45.5 ± 1.3 | 25.2 ± 0.7 | 0.437 ± 0.013 | 0.624 ± 0.010 |
| **Claude 3.5 Sonnet** | 73.66 ± 0.22 | 51.44 ± 0.34 | 53.11 ± 0.24 | 31.2 ± 0.7 | 52.98 ± 0.25 | 30.6 ± 0.5 | 0.5112 ± 0.0026 | 0.616 ± 0.018 |
| **GPT-4o** | 73.4 ± 0.5 | 52.9 ± 0.9 | 50.3 ± 0.9 | 33.0 ± 1.0 | 50.2 ± 0.9 | 32.5 ± 1.0 | 0.485 ± 0.010 | 0.639 ± 0.005 |
| **GPT-4o mini** | 68.5 ± 0.8 | 46.9 ± 0.9 | 43.44 ± 0.25 | 26.9 ± 1.2 | 43.17 ± 0.25 | 26.8 ± 1.2 | 0.4134 ± 0.0027 | 0.567 ± 0.006 |
| **Gemini 1.5 Flash** | 55.0 ± 0.6 | 37.3 ± 0.7 | 34.4 ± 0.9 | 16.9 ± 0.8 | 34.3 ± 0.9 | 16.0 ± 0.7 | 0.321 ± 0.009 | 0.545 ± 0.013 |
| **Gemini 1.5 Pro** | 63.7 ± 0.9 | 44.4 ± 0.8 | 42.6 ± 0.9 | 24.0 ± 1.0 | 42.3 ± 0.8 | 23.4 ± 1.0 | 0.401 ± 0.009 | 0.596 ± 0.007 |
| **Plurality Vote** | 72.0 ± 0.5 | 50.3 ± 1.2 | 48.4 ± 0.8 | 28.9 ± 1.7 | 48.4 ± 0.8 | 28.5 ± 1.7 | 0.465 ± 0.008 | 0.720 ± 0.006 |

**Supplementary Table 2.** LLM performance on only the portion of Tabula Sapiens v2 that was not collected as part of Tabula Sapiens v1.

| | Binary (%) | | Perfect Match (%) | | Extract String Match (%) | | Kappa | |
|---|---|---|---|---|---|---|---|---|
| | Cells | By Cell Type | Cells | By Cell Type | Cells | By Cell Type | With Manual | Average With LLMs |
| **Claude 3 Haiku** | 75.8 ± 2.7 | 66.2 ± 3.5 | 65 ± 7 | 51 ± 11 | 65 ± 7 | 51 ± 11 | 0.62 ± 0.07 | 0.67 ± 0.06 |
| **Claude 3 Opus** | 82.7 ± 2.1 | 68.2 ± 2.6 | 76 ± 5 | 57 ± 9 | 76 ± 5 | 57 ± 9 | 0.73 ± 0.04 | 0.70 ± 0.04 |
| **Claude 3.5 Sonnet** | 84.5 ± 0.8 | 68.0 ± 1.4 | 76.1 ± 2.7 | 55 ± 7 | 76.0 ± 2.9 | 54 ± 7 | 0.734 ± 0.025 | 0.69 ± 0.05 |
| **Command R Plus** | 76.5 ± 3.2 | 61.4 ± 2.5 | 66 ± 7 | 48 ± 12 | 66 ± 7 | 48 ± 12 | 0.63 ± 0.07 | 0.62 ± 0.06 |
| **GPT-4** | 76.6 ± 0.8 | 64.7 ± 2.5 | 66 ± 7 | 47 ± 11 | 66 ± 7 | 47 ± 11 | 0.63 ± 0.07 | 0.67 ± 0.09 |
| **GPT-4o** | 80.7 ± 2.3 | 69.4 ± 2.7 | 71 ± 6 | 55 ± 10 | 71 ± 6 | 55 ± 11 | 0.68 ± 0.06 | 0.71 ± 0.04 |
| **GPT-4o mini** | 73.6 ± 1.8 | 65.1 ± 1.5 | 65 ± 7 | 49 ± 10 | 65 ± 7 | 49 ± 10 | 0.62 ± 0.06 | 0.70 ± 0.05 |
| **Gemini 1.5 Flash** | 66 ± 5 | 59.1 ± 3.3 | 50 ± 7 | 43 ± 10 | 50 ± 7 | 42 ± 11 | 0.45 ± 0.06 | 0.57 ± 0.05 |
| **Gemini 1.5 Pro** | 78.1 ± 2.5 | 64.8 ± 2.3 | 68 ± 6 | 50 ± 9 | 68 ± 6 | 50 ± 9 | 0.65 ± 0.06 | 0.65 ± 0.05 |
| **Llama 3.1 405B Instruct** | 81 ± 4 | 66 ± 4 | 68 ± 6 | 50 ± 11 | 68 ± 6 | 50 ± 11 | 0.65 ± 0.06 | 0.65 ± 0.05 |
| **Llama 3.1 70B Instruct** | 69 ± 4 | 60.6 ± 2.9 | 61 ± 4 | 47 ± 10 | 61 ± 4 | 47 ± 10 | 0.576 ± 0.033 | 0.64 ± 0.04 |
| **Llama 3.1 8B Instruct** | 54 ± 9 | 53 ± 6 | 45 ± 8 | 42 ± 11 | 45 ± 8 | 42 ± 11 | 0.41 ± 0.06 | 0.50 ± 0.07 |
| **Mistral Large** | 75.4 ± 2.6 | 65.0 ± 3.4 | 63 ± 5 | 51 ± 10 | 63 ± 5 | 50 ± 10 | 0.59 ± 0.04 | 0.68 ± 0.04 |
| **Plurality Vote** | 78.6 ± 1.8 | 68.0 ± 2.4 | 72 ± 5 | 55 ± 9 | 72 ± 5 | 55 ± 9 | 0.70 ± 0.04 | 0.76 ± 0.04 |

**Supplementary Table 3.** LLM performance when post-processed by GPT-4o.

| | Binary (%) | | Perfect Match (%) | | Extract String Match (%) | | Kappa | |
|---|---|---|---|---|---|---|---|---|
| | **Cells** | **By Cell Type** | **Cells** | **By Cell Type** | **Cells** | **By Cell Type** | **With Manual** | **Average With LLMs** |
| **Claude 3 Haiku** | 78.42 ± 0.29 | 66.2 ± 1.0 | 58.9 ± 1.3 | 36.3 ± 2.8 | 58.5 ± 1.6 | 35.2 ± 3.1 | 0.557 ± 0.017 | 0.620 ± 0.007 |
| **Claude 3 Opus** | 82.09 ± 0.22 | 68.3 ± 0.7 | 68.7 ± 0.9 | 44.8 ± 2.0 | 68.2 ± 1.4 | 43.5 ± 2.5 | 0.658 ± 0.014 | 0.677 ± 0.012 |
| **Claude 3.5 Sonnet** | 83.0 ± 0.6 | 68.6 ± 0.7 | 70.4 ± 1.1 | 45.2 ± 2.4 | 69.9 ± 1.5 | 43.6 ± 3.5 | 0.678 ± 0.016 | 0.653 ± 0.016 |
| **Command R Plus** | 76.3 ± 0.4 | 57.4 ± 1.1 | 61.3 ± 0.8 | 34.3 ± 2.0 | 60.4 ± 1.3 | 32.8 ± 3.0 | 0.575 ± 0.014 | 0.604 ± 0.014 |
| **GPT-4** | 79.4 ± 0.7 | 64.0 ± 0.7 | 57.9 ± 1.7 | 34.2 ± 2.7 | 57.2 ± 1.8 | 33.2 ± 3.5 | 0.542 ± 0.019 | 0.591 ± 0.024 |
| **GPT-4o** | 80.4 ± 0.5 | 66.8 ± 1.1 | 67.3 ± 0.6 | 45.0 ± 2.7 | 66.8 ± 1.0 | 43.5 ± 3.3 | 0.644 ± 0.011 | 0.691 ± 0.013 |
| **GPT-4o mini** | 76.13 ± 0.33 | 64.2 ± 1.3 | 60.2 ± 1.0 | 37.3 ± 1.9 | 59.9 ± 1.2 | 36.4 ± 2.4 | 0.571 ± 0.012 | 0.649 ± 0.014 |
| **Gemini 1.5 Flash** | 67.90 ± 0.30 | 58.8 ± 0.4 | 47.9 ± 0.5 | 32.5 ± 2.2 | 47.5 ± 0.7 | 30.2 ± 3.1 | 0.447 ± 0.007 | 0.529 ± 0.014 |
| **Gemini 1.5 Pro** | 79.3 ± 1.5 | 66.9 ± 1.1 | 63.1 ± 2.3 | 40.2 ± 3.0 | 62.7 ± 2.7 | 39 ± 4 | 0.602 ± 0.028 | 0.628 ± 0.021 |
| **Llama 3.1 405B Instruct** | 82.3 ± 1.2 | 66.0 ± 1.4 | 65.6 ± 0.5 | 36.8 ± 2.7 | 65.2 ± 0.4 | 35.6 ± 3.1 | 0.625 ± 0.005 | 0.659 ± 0.011 |
| **Llama 3.1 70B Instruct** | 72.6 ± 1.8 | 59.8 ± 2.2 | 58.7 ± 2.0 | 35.8 ± 2.0 | 58.3 ± 1.8 | 35.0 ± 2.5 | 0.557 ± 0.019 | 0.623 ± 0.015 |
| **Llama 3.1 8B Instruct** | 56.0 ± 2.1 | 47.5 ± 1.5 | 43.6 ± 2.1 | 27.3 ± 1.8 | 42.9 ± 2.0 | 25.3 ± 2.7 | 0.397 ± 0.020 | 0.488 ± 0.019 |
| **Mistral Large** | 77.56 ± 0.33 | 62.9 ± 1.3 | 61.4 ± 1.5 | 39.9 ± 2.2 | 61.0 ± 1.7 | 38.3 ± 3.0 | 0.587 ± 0.018 | 0.655 ± 0.010 |
| **Plurality Vote** | 80.47 ± 0.26 | 67.4 ± 1.3 | 69.2 ± 0.5 | 45.5 ± 1.8 | 68.8 ± 0.6 | 44.2 ± 2.5 | 0.665 ± 0.006 | 0.742 ± 0.009 |

**Supplementary Table 4.** Comparison of agreement metrics calculated using two different LLMs for post-processing of annotations (Claude 3.5 Sonnet and GPT-4o).

| | Pearson's R | P |
|---|---|---|
| **Overall Binary (% of Cells)** | 0.99 | 6.5e-13 |
| **Overall Binary (% of Cell Types)** | 0.97 | 9.2e-09 |
| **Perfect Match (% of Cells)** | 0.99 | 9.5e-12 |
| **Perfect Match (% of Cell Types)** | 0.98 | 3.0e-09 |
| **Exact String Match (% of Cells)** | 0.99 | 1.5e-11 |
| **Exact String Match (% of Cell Types)** | 0.98 | 7.1e-10 |
| **Categorical Agreement (% of Cells)_1.0** | 0.99 | 9.5e-12 |
| **Categorical Agreement (% of Cells)_0.5** | 0.94 | 7.1e-07 |
| **Categorical Agreement (% of Cells)_0.0** | 1.00 | 2.0e-16 |
| **Categorical Agreement (% of Cell Types)_1.0** | 0.98 | 3.0e-09 |
| **Categorical Agreement (% of Cell Types)_0.5** | 0.78 | 9.9e-04 |
| **Categorical Agreement (% of Cell Types)_0.0** | 0.98 | 7.0e-10 |
| **Kappa with Manual Annotations** | 0.99 | 1.9e-11 |
| **Average Kappa with Other LLMs** | 0.99 | 8.5e-12 |

**Supplementary Table 5.** LLM performance when rated by GPT-4o.

| | Binary (%) | | Perfect Match (%) | | Extract String Match (%) | | Kappa | |
|---|---|---|---|---|---|---|---|---|
| | Cells | By Cell Type | Cells | By Cell Type | Cells | By Cell Type | With Manual | Average With LLMs |
| **Claude 3 Haiku** | 80.4 ± 1.4 | 71.2 ± 2.1 | 61.8 ± 2.3 | 47 ± 4 | 61.8 ± 2.3 | 47 ± 4 | 0.589 ± 0.023 | 0.652 ± 0.023 |
| **Claude 3 Opus** | 84.1 ± 1.9 | 72.5 ± 2.0 | 72.8 ± 2.6 | 54 ± 4 | 72.7 ± 2.6 | 54 ± 4 | 0.704 ± 0.026 | 0.711 ± 0.020 |
| **Claude 3.5 Sonnet** | 87.4 ± 2.9 | 74.9 ± 1.9 | 74.4 ± 2.7 | 54 ± 4 | 74.3 ± 2.6 | 53 ± 4 | 0.721 ± 0.027 | 0.697 ± 0.026 |
| **Command R Plus** | 78.0 ± 1.3 | 59.3 ± 3.4 | 64.5 ± 2.6 | 40 ± 5 | 64.5 ± 2.6 | 40 ± 5 | 0.616 ± 0.027 | 0.646 ± 0.026 |
| **GPT-4** | 80.3 ± 1.8 | 68.8 ± 2.3 | 64 ± 4 | 44 ± 5 | 64 ± 4 | 44 ± 5 | 0.61 ± 0.05 | 0.65 ± 0.04 |
| **GPT-4o** | 83.9 ± 2.6 | 73.7 ± 3.3 | 70.4 ± 2.5 | 54 ± 5 | 70.4 ± 2.5 | 54 ± 6 | 0.680 ± 0.026 | 0.721 ± 0.021 |
| **GPT-4o mini** | 80.4 ± 2.6 | 69.5 ± 2.1 | 63.4 ± 3.0 | 47 ± 6 | 63.4 ± 3.0 | 47 ± 6 | 0.605 ± 0.031 | 0.681 ± 0.022 |
| **Gemini 1.5 Flash** | 73.1 ± 2.8 | 65.7 ± 2.9 | 51.0 ± 2.5 | 41 ± 5 | 51.0 ± 2.5 | 41 ± 5 | 0.478 ± 0.024 | 0.561 ± 0.020 |
| **Gemini 1.5 Pro** | 80.5 ± 2.2 | 72.1 ± 1.9 | 65.1 ± 2.4 | 50 ± 5 | 65.1 ± 2.4 | 50 ± 5 | 0.625 ± 0.024 | 0.658 ± 0.019 |
| **Llama 3.1 405B Instruct** | 83.6 ± 1.3 | 70.0 ± 3.2 | 69.5 ± 2.6 | 47 ± 5 | 69.3 ± 2.6 | 47 ± 5 | 0.667 ± 0.027 | 0.690 ± 0.021 |
| **Llama 3.1 70B Instruct** | 77.5 ± 2.6 | 67.3 ± 2.1 | 64 ± 4 | 45 ± 4 | 64 ± 4 | 45 ± 5 | 0.62 ± 0.04 | 0.665 ± 0.022 |
| **Llama 3.1 8B Instruct** | 63.2 ± 3.4 | 58 ± 4 | 47.7 ± 3.2 | 37 ± 5 | 47.6 ± 3.2 | 36 ± 6 | 0.440 ± 0.031 | 0.526 ± 0.030 |
| **Mistral Large** | 81.4 ± 1.9 | 70.1 ± 2.6 | 64.9 ± 2.7 | 50 ± 5 | 64.8 ± 2.8 | 49 ± 5 | 0.623 ± 0.027 | 0.696 ± 0.024 |
| **Plurality Vote** | 83.5 ± 2.3 | 73.1 ± 2.5 | 72.4 ± 2.1 | 55 ± 5 | 72.3 ± 2.1 | 55 ± 5 | 0.700 ± 0.022 | 0.770 ± 0.018 |

**Supplemental Table 6.** Comparison of agreement metrics calculated using two different LLMs (Claude 3.5 Sonnet and GPT-4o) to rate annotations.

| | Pearson's R | P |
|---|---|---|
| **Overall Binary (% of Cells)** | 0.99 | 6.2e-11 |
| **Overall Binary (% of Cell Types)** | 0.96 | 6.9e-08 |
| **Perfect Match (% of Cells)** | 1.00 | 2.6e-37 |
| **Perfect Match (% of Cell Types)** | 1.00 | 5.4e-22 |
| **Exact String Match (% of Cells)** | 1.00 | 0.0e+00 |
| **Exact String Match (% of Cell Types)** | 1.00 | 0.0e+00 |
| **Categorical Agreement (% of Cells)_1.0** | 1.00 | 2.6e-37 |
| **Categorical Agreement (% of Cells)_0.5** | 0.99 | 7.6e-11 |
| **Categorical Agreement (% of Cells)_0.0** | 0.99 | 9.4e-13 |
| **Categorical Agreement (% of Cell Types)_1.0** | 1.00 | 5.4e-22 |
| **Categorical Agreement (% of Cell Types)_0.5** | 0.95 | 1.8e-07 |
| **Categorical Agreement (% of Cell Types)_0.0** | 0.97 | 1.7e-08 |
| **Kappa with Manual Annotations** | 1.00 | 0.0e+00 |
| **Average Kappa with Other LLMs** | 1.00 | 0.0e+00 |

**Supplementary Table 7.** LLM performance (Claude 3.5 Sonnet) during prompt ablation.

| | Binary (%) | | Perfect Match (%) | | Extract String Match (%) | | Kappa |
|---|---|---|---|---|---|---|---|
| | Cells | By Cell Type | Cells | By Cell Type | Cells | By Cell Type | With Manual |
| **Unablated** | 78 ± 4 | 66.1 ± 3.4 | 47 ± 17 | 30 ± 15 | 41 ± 21 | 26 ± 19 | 0.39 ± 0.21 |
| **Base prompt detuned** | — | — | — | — | — | — | — |
| **No tissue context** | 81.5 ± 2.2 | 59.3 ± 3.3 | 46 ± 14 | 21 ± 14 | 44 ± 16 | 20 ± 15 | 0.42 ± 0.16 |
| **No system prompt** | 78 ± 5 | 61 ± 7 | 42 ± 24 | 30 ± 17 | 38 ± 28 | 25 ± 21 | 0.36 ± 0.28 |
| **Randomize gene list order** | 80.2 ± 2.3 | 63 ± 4 | 48 ± 14 | 28 ± 15 | 42 ± 19 | 24 ± 18 | 0.40 ± 0.19 |

**Supplementary Table 8.** Example biological process annotations of known gene lists using an LLM (Claude 3.5 Sonnet).

| GO Biological Process Term | Genes | LLM annotation |
|---|---|---|
| 'De Novo' AMP Biosynthetic Process (GO:0044208) | ATIC, PAICS, PFAS, ADSS1, ADSS2, GART | Purine biosynthesis pathway |
| 'De Novo' Post-Translational Protein Folding (GO:0051084) | SDF2L1, HSPA9, CCT2, HSPA6, ST13, ENTPD5, HSPA1L, HSPA5, PTGES3, HSPA8, HSPA7, DNAJB13, HSPA2, DNAJB14, HSPE1, DNAJC18, GAK, DNAJC7, DNAJB12, HSPA1A, ST13P5, HSPA1B, ERO1A, SELENOF, HSPA14, HSPA13, DNAJB1, CHCHD4, DNAJB5, DNAJB4, SDF2, UGGT1 | Protein folding and chaperone-mediated quality control. |
| 2-Oxoglutarate Metabolic Process (GO:0006103) | IDH1, PHYH, GOT2, MRPS36, GOT1, IDH2, ADHFE1, GPT2, TAT, DLST, OGDHL, L2HGDH, D2HGDH, OGDH | Mitochondrial tricarboxylic acid (TCA) cycle and related metabolic pathways. |
